# SlideCNA: Spatial copy number alteration detection from Slide-seq-like spatial transcriptomics data

**DOI:** 10.1101/2022.11.25.517982

**Authors:** Diane Zhang, Asa Segerstolpe, Michal Slyper, Julia Waldman, Evan Murray, Ofir Cohen, Orr Ashenberg, Daniel Abravanel, Judit Jané-Valbuena, Simon Mages, Ana Lako, Karla Helvie, Orit Rozenblatt-Rosen, Scott Rodig, Fei Chen, Nikhil Wagle, Aviv Regev, Johanna Klughammer

## Abstract

Solid tumors are spatially heterogeneous in their genetic, molecular and cellular composition, and this variation can be meaningful for diagnosis, prognosis and therapy. Recent spatial profiling studies have mostly charted genetic and RNA variation in tumors separately. To leverage the potential of RNA to identify copy number alterations (CNAs), we developed SlideCNA, a computational tool to extract sparse spatial CNA signals from spatial transcriptomics data, using expression-aware spatial binning. We test SlideCNA on simulated and real Slide-seq data of metastatic breast cancer (MBC) and demonstrate its potential for spatial sub-clone detection.

## Background

The spatial organization of tumors at the genetic, expression, and histological levels provides important insights into a tumor’s evolution, growth, microenvironment, and response to therapy (1). In spot-based sequencing methods, such as Spatial Transcriptomics (ST, commercialized as Visium) (2) and Slide-Seq (3), tissue is positioned on a glass slide coated with RNA capture probes or beads, such that RNA molecules are spatially barcoded in an untargeted, arrayed fashion. Recent advances have led to increasingly higher resolution (55μm in Visium-ST and 10μm in Slide-seq), albeit at the cost of higher sparsity (4,5). ST studies have provided insights into the cellular and molecular organization of tumor tissue ecosystems, but – aside from direct experimental measurements (6) – these have not yet been associated with the genetic state of malignant cells, including tumor cell clonality. Relating the spatial context of clonal genetic events to cell states in the tumor ecosystem can help distinguish malignant and non-malignant cells and shed light on the impact of the microenvironment on the mutational landscape, therapeutic resistance, progression, and metastasis.

In previous studies, we developed InferCNV, a computational method that uses single-cell RNA-seq (scRNA-seq) profiles in cancer cells to infer genomic copy number alterations (CNAs) (7), and it has since been applied to spatial transcriptomics of tumor tissues for spatial CNA detection (8). An additional method, STARCH, further utilizes spatial information to call CNAs and infer clones (9). However, despite these advancements, the relatively low spatial resolution of ST methods, where each measurement consists of several cells (or fractions of cells), continues to pose a challenge, as it mixes signals of different clones and cell types.

## Results and Discussion

To address this challenge, we developed SlideCNA to improve CNA inference and clone calling in spatial transcriptomics data by leveraging the increased resolution of Slide-seq data with a spatio-molecular binning step to address signal sparsity (**Fig. 1a, Methods**). Similar to InferCNV, SlideCNA implements an expression-based smoothing approach across each chromosome with a pyramidal weighting average scheme. Then, SlideCNA adjusts all expression values by the average expression of a user-defined set of reference beads (or spots) for each gene and centers these values to produce relative smoothed expression intensities. To overcome Slide-seq’s sparsity, SlideCNA uses a spatial binning step to combine neighboring beads into bins that increase the signal (counts), while maintaining spatial structure. To this end, SlideCNV computes bead-by-bead distances in both expression and physical space, then takes a weighted linear combination of the expression and spatial distance matrices and hierarchically clusters this combined pseudodistance matrix, to group beads with similar expression profiles that are also proximal in physical space. SlideCNA partitions the beads into bins with a user-defined maximum number of beads per bin, calculates bin expression intensities as an average across the constituent beads, and normalizes and scales these intensities for UMI count, to generate CNA scores.

**Fig. 1.**
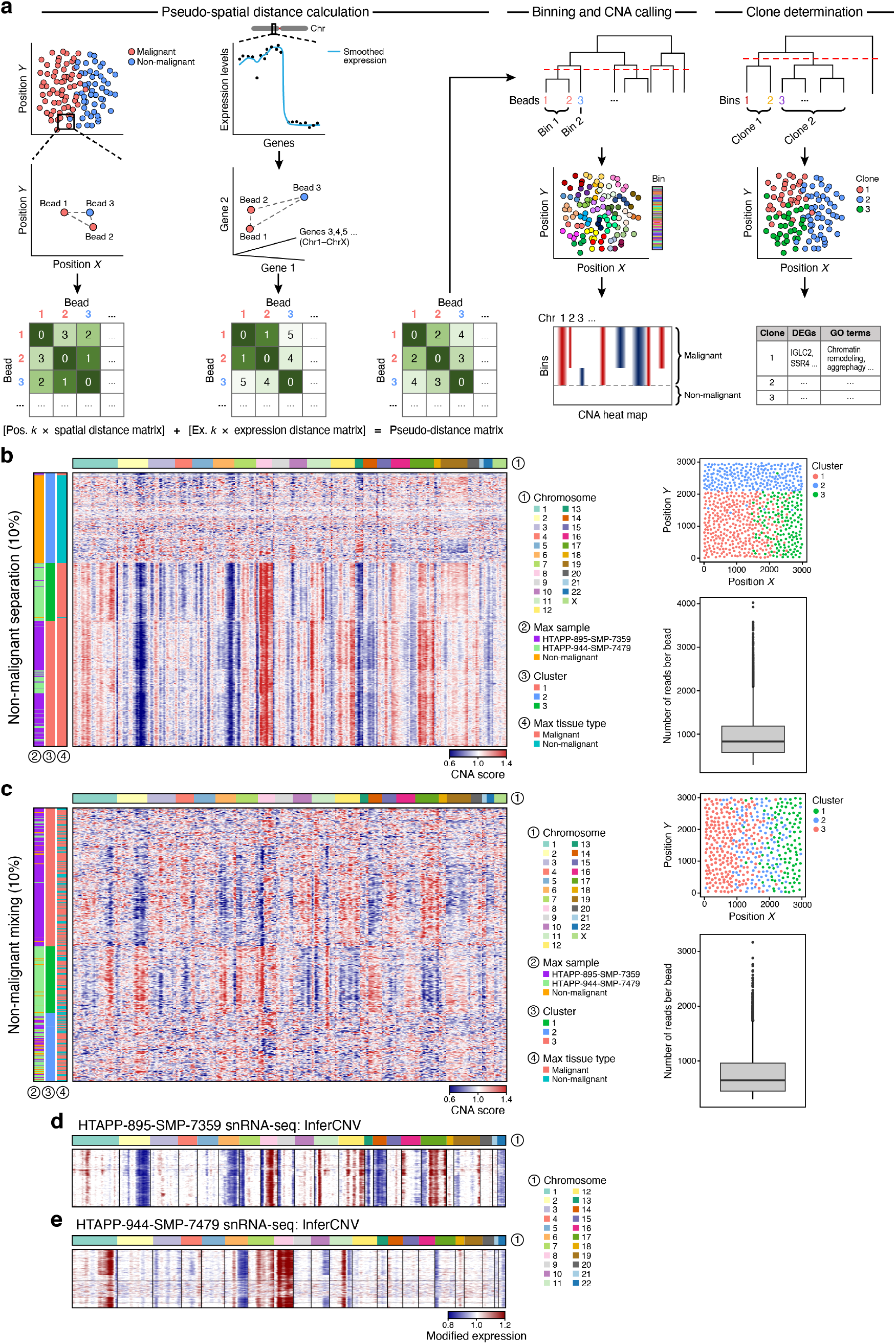
SlideCNA methodology schematic and *in silico* spatial results. **a**, SlideCNA methodology involves calculating bead distances by a combined spatial- and expression-based pseudo-distance, binning beads based on pseudo-distance, and determining malignant clones based on CNA profiles. **b,c,** SlideCNA heat map (amplification > 1, deletion < 1), spatial plot of bins colored by assigned cluster, and boxplot of number of reads per bin after filtering for beads with >300 counts across all genes for the *in silico* non-malignant-separated dataset **(b)** and non-malignant-mixed dataset **(c)** with counts downsampled to 10%. **d,e,** Summary of the InferCNV CNA profiles of malignant nuclei for snRNA-seq data of HTAPP-895-SMP-7359 **(d)** and HTAPP-944-SMP-7479 **(e)**.

We first tested SlideCNA’s ability to recover a ground truth CNA profile using an *in silico* (simulated) Slide-Seq dataset with a known clonal structure, which we constructed from two independent single-nucleus RNA-seq (snRNA-Seq) datasets of metastatic breast cancer (MBC) (**Supplementary Fig. 1a**). Mimicking the process of RNA capture on Slide-seq beads for a tissue section, we placed 20,000 nucleus profiles on a 2-dimensional square, positioning malignant cells of each sample on opposite regions in a gradient-like manner, such that cells of both samples would be equally mixed in the middle (**Supplementary Fig. 1b**, bottom label). Non-malignant cell type nucleus profiles were either positioned (1) separately but adjacent to the malignant nucleus profiles, reflecting a discrete compartmentalized normal population (**Supplementary Fig. 1b**, left and **1c**), or (2) randomly across the square, reflecting a more challenging intermixing of non-malignant and malignant cells (*e.g*., due to infiltration) (**Supplementary Fig. 1b**, right, and **1d**). For each scenario, we applied a Gaussian kernel to create a square of 10,000 beads with combined expression values of the constituent nuclei profiles (or their fractions) (**Supplementary Fig. 1b, bottom**). To roughly approximate the sparsity of Slide-seq data, we downsampled the *in silico* beads to 10%.

Applied to these *in silico* datasets, SlideCNA generated CNA profiles with unique patterns in two malignant clusters, recapitulating the original positioning of sample nucleus profiles, both in the normal separation and normal mixing case (**Fig. 1b-e, Supplementary Fig. 1e,f**). In the normal separation and mixing cases, we observed consistent results without downsampling, and at 25% and 10% downsampling, but not with 2% downsampling (**Supplementary Figs. 2 and 3**), where 10% downsampling corresponds to the typical UMI counts per bead in real Slide-seq data. This suggests that SlideCNA can potentially recover subclones with distinct CNA profiles in Slide-seq-like data.

Next, we applied SlideCNA to two real Slide-seq samples from metastatic breast cancer tumors with matching snRNA-seq. In the first tumor (HTAPP-895-SMP-7359), we annotated non-malignant beads as a reference against which contiguous chromosomal expression changes could be scored as malignant and to infer CNAs. Chromosome-wide CNA scores were spatially heterogeneous (**Fig. 2a,b**). Chromosomes 13, 21, and 22 showed strong CNA signals (deletion for chromosomes 13 and 22 and amplification for chromosome 21), specifically in regions annotated as malignant, while chromosomes 11 and 12 consistently did not show CNAs across space. Malignant beads partitioned into two clusters that mapped to different regions by their inferred CNA profile, with few significant differentially expressed genes between clusters (**Fig. 2c-e,** clusters 1 and 2). Non-malignant beads occupied their own spatially distinct cluster as well (**Fig. 2c**, cluster 3). In the second tumor (HTAPP-944-SMP-7479), since there was no clear zonation of cell types and hence no clearly delineated non-malignant reference beads, we used the non-malignant beads from HTAPP-895-SMP-7359 as reference (**Methods**). From this, we recovered a single malignant bead cluster with distinct CNA patterns from the non-malignant reference population, which partitioned into two clusters, and detected only slight spatial variability in CNAs on chromosomes 8, 21, and 22 (**Fig. 2f-j**).

**Fig. 2.**
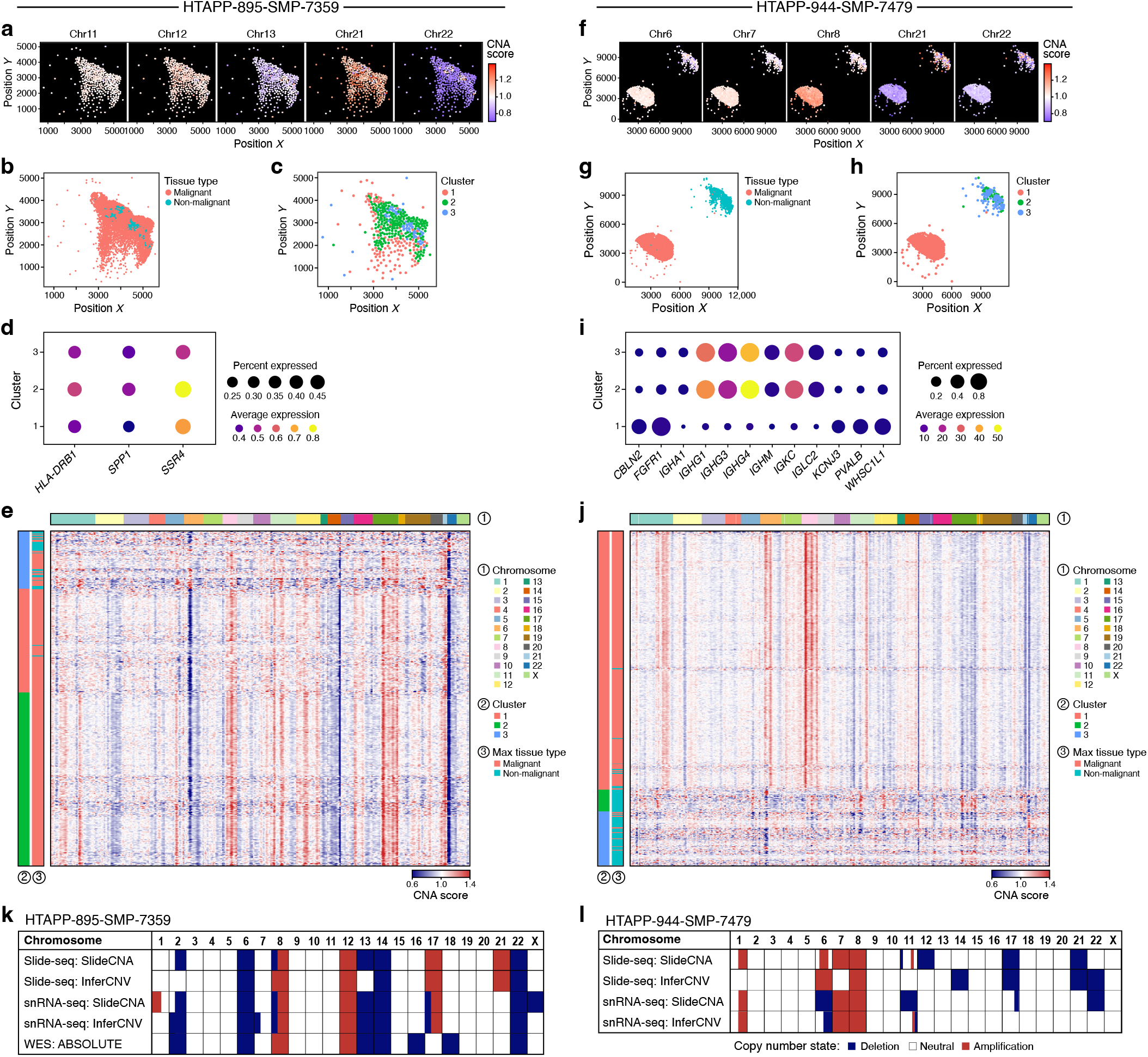
SlideCNA identifies spatial patterns in Slide-seq MBC samples. **a,f**, Beads were colored by mean CNA score across all genes per chromosome for chrs. 11, 12, 13, 21, and 22 (HTAPP-895-SMP-7359) or chrs. 6,7,8,21, and 22 (HTAPP-944-SMP-7479) on spatial axes. **b,g,** Slide-seq beads were annotated as non-malignant (blue) or malignant (pink) after *de novo* reference bead selection. HTAPP-944-SMP-7479 contains reference beads transplanted from the HTAPP-895-SMP-7359 sample and spatially offset in the upper right corner by (100 + max x coordinate of HTAPP-944-SMP-7479, 100 + max y coordinate of HTAPP-944-SMP-7479) (**g**). **c,h,** Binned beads were colored by SlideCNA-defined cluster designation on spatial axes. **d,i,** Top DEGs for each malignant cluster detected from the SlideCNA profile. DEGs were colored by average cluster expression and sized by the percent of beads expressing that gene in the cluster. **e,j,** SlideCNA heat map of malignant and non-malignant binned beads annotated with cluster assignment. (Note that some non-malignant beads have an artifactual call inverse to the malignant beads, such as the non-malignant ‘amplification’ where there is a malignant deletion on chromosome 17.) **a-e** refer to sample HTAPP-895-SMP-7359 and **f-j** refer to sample HTAPP-944-SMP-7479. **k,l,** Comparison of manually assigned SlideCNA and Infercnv calls for both Slide-seq and snRNA-seq data and ABSOLUTE calls for WES data (WES data for **k**, HTAPP-895-SMP-7359, only) for HTAPP-895-SMP-7359 (**k**) and HTAPP-944-SMP-7479 (**l**).

The SlideCNA-inferred profiles were consistent with those obtained from the matching snRNA-seq called with either InferCNV or SlideCNA with non-spatial parameters. As further genomic validation, for HTAPP-895-SMP-7359, we also had access to matching whole exome sequencing (WES) data, and obtained CNA profiles with ABSOLUTE (10). To compare profiles, we visually assessed the CNA profiles from each method, and manually assigned copy number states to each chromosome for each sample and method (**Fig. 2k,l**). Slide-seq CNA profiles were largely consistent with those from snRNA-seq, both using InferCNV (current gold standard developed for scRNA-seq) and SlideCNA applied on the snRNA-seq data without spatial parameters (**Supplementary Fig. 1e,f and 4a-e**). When comparing SlideCNA and InferCNV’s calls on Slide-seq data, SlideCNA yielded CNAs at higher chromosomal resolution, including several arm-level CNAs detected only by SlideCNA, but not InferCNV, which were also present in the snRNA-seq calls, such as CNAs on chromosomes 2q, 8p, and 13 for HTAPP-895-SMP-7359 and chromosome 1q for HTAPP-944-SMP-7479 (although InferCNV may have inferred Chromosome 8p more accurately based on WES rather than snRNA-seq **(Fig. 2k,l and Supplementary Fig. 4a-d**). This highlights SlideCNA’s ability to preserve the spatial landscape of CNAs, while overcoming signal sparsity. Furthermore, for HTAPP-895-SMP-7359, copy number profiles from both snRNA-seq and Slide-seq data were largely consistent with those of ABSOLUTE analysis on WES, further supporting their accuracy, though in some cases there were discrepancies, including 8p (**Fig. 2k** and **Supplementary Fig. 4e**). Most of SlideCNA’s major CNA calls on Slide-seq data were consistent with ABSOLUTE’s WES CNA patterns for HTAPP-895-SMP-7359, except for the lack of chr16 and 18 deletions and uniquely detected chr17 and 21 amplifications (**Supplementary Fig. 4e**).

Finally, we compared SlideCNA’s calls on both binned Slide-seq samples with those of STARCH, the existing spatially-aware and cluster-assigning CNA calling tool (9). When setting the number of malignant bead clusters to two, STARCH inferred a large number of amplifications for HTAPP-895-SMP-7359 and of deletions for HTAPP-944-7479 (**Supplementary Fig. 5a-d)**. We conclude that these were not reliable because they vastly differed from CNA profiles of SlideCNA and InferCNV for both Slide-seq and snRNA-seq data, and from ABSOLUTE’s predictions on WES. Due to the poor CNA calls, spatial interpretation of STARCH-assigned clusters was limited. This underscores the unique challenges that Slide-seq data present in CNA calling.

## Conclusion

We demonstrated that SlideCNA simultaneously captures spatial signal and identifies CNAs from Slide-seq data, with potential power to detect subclonal structure, despite limited capture efficiency and imperfect separation between normal and malignant expression. SlideCNA should help empower studies of tumor ecosystems in large-scale spatial CNA screening and expression analysis in clinical samples.

## Methods

### Tissue samples

Matching snRNA-seq and Slide-seq data were generated for two breast cancer liver metastases (HTAPP-895-SMP-7359 and HTAPP-944-SMP-7479) of two different patients **(Fig. 1a)**. Tissues were collected as described previously (11). Specifically, MBC OCT-frozen samples were obtained from the Center for Cancer Precision Medicine Bank. Fresh samples (core-needle biopsies) were frozen in optimal cutting temperature compound (OCT, Tissue-Tek Sakura). Cores were pre-coated with OCT by putting a thin layer of OCT down in the cryomold before placing an individual core in the center of the OCT mold in a straight line and adding additional OCT to fill the cryomold. The cryomold was then placed on dry ice for 5-15 minutes until the block was opaque before storing it at −80°C.

### Single nucleus RNA-seq

SnRNA-seq was performed as described previously (11). Specifically, frozen tissue was placed on ice and in one well of a plate (Stem Cell Technologies, cat. no. 38015) and 1ml of TST buffer was added to the well. Tissue was kept on ice and cut into pieces with Noyes Spring Scissors (Fine Science Tools, cat. no. 15514-12) for 10 min. Tissue mixture was filtered through a 40 μm Falcon cell strainer (ThermoFisher Scientific, cat. no. 08-771-1). The well was washed, filtered with 1 ml of detergent buffer solution, and 3 ml of 1× ST buffer were added to a total well volume of 5 ml. The solution was centrifuged in a 15 ml Eppendorf tube for 5 min at 500*g* and 4 °C in a swinging bucket centrifuge. Pellet was resuspended in 1× ST buffer, with a resuspension volume of 100-200 μl based on pellet size. The single nucleus suspension was filtered through a 35 μm Falcon cell strainer (Corning, cat. no. 352235). 10,000 nuclei were selected with a C-chip disposable hemocytometer (VWR, cat. no. 82030-468) and transferred to Chromium chips for the Chromium Single Cell 3’ Library (V3, PN-1000075) per manufacturer’s instructions (10x Genomics).

### Slide-Seq

Slide-seq was performed as previously described (5). Specifically, core biopsies embedded in OCT (Tissue-Tek Sakura) and kept at −80°C were sectioned at 10μm thickness onto one Slide-seq puck each. Loaded Slide-seq pucks were incubated with hybridization buffer (6x SSC with 2 U/μl RNase inhibitor (Lucigen, 30281)) and subjected to first-strand cDNA synthesis (1x Maxima RT buffer, 1 mM of each dNTP, 0.05U/μl RNase inhibitor (Lucigen, 30281), 2.5uM Template switch oligo (TSO) and 10U/μl Maxima H Minus reverse transcriptase (ThermoFisher, EP0742) followed by tissue digestion (200 mM Tris-Cl pH 7.5, 400 mM NaCl, 4% SDS, 10 mM EDTA with 1:50 proteinase K (New England BioLabs, P8107S)), as previously described. Puck beads were released into suspension and washed in wash buffer (10 mM Tris pH 8, 1 mM EDTA, 0.01% Tween-20) before incubation with exonuclease I (1x ExoI buffer with 10U/μl Exonuclease I (New England BioLabs, M0293L)) to prepare for second-strand cDNA synthesis which was performed using the following reaction mix: 1x Maxima RT buffer, 1 mM of each dNTP, 10 μM dN-SMRT oligonucleotide, and 0.125U/μl Klenow enzyme (NEB, M0210). The sample was subjected to PCR amplification with 1x Terra Direct PCR mix buffer, 2 μl Terra polymerase (Takara, 639270), 2 μM TruSeq PCR handle primer, and 2 μM SMART PCR primer (98°C for 2 min; four cycles of 98°C for 20 s, 65°C for 45 s and 72°C for 3 min; 11 cycles of 98°C for 20 s, 67°C for 20 s and 72°C for 3 min; 72°C for 5 min; hold at 4°C). cDNA was purified using AMPure XP beads and quantified on a Bioanalyzer High Sensitivity DNA chip (Agilent, 5067-4626) and on a Qubit high sensitivity dsDNA kit (Invitrogen, Q32851). 600 pg of cDNA were tagmented with a Nextera XT kit (Illumina, FC-131-1096) and libraries were indexed with TruSeq5 and the N700 series barcoded index primers. Libraries were purified using AMPure XP beads and sequenced on an Illumina NextSeq High Output flow cell with the settings: read1 44 bases, read2 39 bases, and index1 8 bases.

### snRNA-Seq data pre-processing

Raw counts matrices were generated from fastq files using the 10x CellRanger pipeline as implemented in Cumulus (12), set to include reads from intronic regions and using the hg38 reference genome together with the corresponding gene annotation. SnRNA-seq raw count data were filtered and genes with ≤ 300 counts across all nuclei were removed. The filtered count matrix was transformed into a Seurat V4 (13) object through CreateSeuratObject, which involves calculating the percentage of mitochondrial transcript contamination with PercentageFeatureSet (default parameters), natural log normalization with a scale factor of 10,000 with NormalizeData, identifying variable genes with FindVariableFeatures (default parameters), scaling normalized counts while regressing out the number of RNA counts and percentage of mitochondrial genes with ScaleData, running PCA with 50 principal components with RunPCA, calculating *k*-nearest neighbors (KNN) with *k*=20 neighbors to construct a shared nearest neighbors (SNN) graph with FindNeighbors, and identifying clusters based on the SNN with FindClusters (default parameters) (14,15).

### Slide-Seq data pre-processing

Counts matrices and bead positions were generated using the Slide-seq pipeline (https://github.com/MacoskoLab/slideseq-tools) and the hg19 reference genome together with the corresponding gene annotation. Slide-seq raw count data were filtered and genes with ≤ 300 counts across all beads were removed. The filtered Slide-seq count matrix and metadata were processed into a Seurat v4 object with the same steps described for snRNA-Seq (above), which was then used to designate beads as normal or malignant (below) (3) **(Supplementary Fig. 6a**).

### *De-novo* reference (non-malignant) cell selection in snRNA-Seq

SingleR v.1.8.0 (16) was used to annotate nucleus profiles and the endothelial, macrophage, natural killer (NK), and monocyte cells were selected as the non-malignant reference cells. **(Supplementary Fig. 6b**).

### *De-novo* reference bead selection in Slide-Seq

The top 100 differentially expressed genes between one bead cluster and all other bead clusters were defined for each Seurat bead cluster and tested for enrichment in Gene Ontology (GO) 2018 Biological Processes with Enrichr (14,15,17).

For HTAPP-895-SMP-7359, each bead cluster was manually annotated as “non-malignant (normal)” or “malignant” by the top 100 differentially expressed genes and enriched GO categories. Because beads (and bead clusters) may have signal from both malignant and non-malignant cells, a malignant marker gene set was defined as the top 50 differentially expressed genes between the one top malignant bead cluster and the one top non-malignant bead cluster (where top clusters were chosen as those with the most malignant/normal-related DEGs and GO terms). A bead’s malignant score was defined as the sum of its gene expression across the malignant gene set using Seurat v4. Beads with a score ≥ 0.1 were assigned as malignant and those < 0.1 as normal.

For HTAPP-944-SMP-7479, where most beads were annotated as malignant, reference (non-malignant) beads were selected from HTAPP-895-SMP-7359 as those in Seurat clusters annotated as immune-related (i.e., non-malignant) modules. Downstream CNA analysis should not be substantially affected by this, because expression patterns are typically similar in non-malignant cells between samples (7). To avoid spatial overlapping of the reference beads, HTAPP-895-SMP-7359 beads were transplanted to occupy a distinct region in the HTAPP-944-SMP-7479 sample space offset by (100 + max x coordinate of HTAPP-944-SMP-7479, 100 + max y coordinate of HTAPP-944-SMP-7479) (**Fig. 2f**).

### SlideCNA

To transform raw counts into gene expression intensities that reflect copy number changes, a modified version of the InferCNV processing pipeline (7) was employed. Beads with ≤ 300 counts across all genes and genes with ≤ 50 counts across all beads were removed, followed by log_2_(TPM+1) transformation of the raw counts matrix **(Supplementary Fig. 6c**). Each gene was centered by subtracting its mean log_2_ expression in the reference beads. Relative expression intensities were capped at ≤ 3 and ≥ −3 and mitochondrial and Y chromosome genes were removed.

Genes were ordered by chromosome, and the centered expression intensities for each bead were smoothed across genomic regions in each chromosome by a pyramidal weighted moving average (default window of *k* = 101 or *k* = *n* if *n* < 101, with *n* = number of genes on the chromosome), with higher weights corresponding to closer genes. Thus, the smoothed expression (*X_i_*) of each gene at position i (*g_i_*) was calculated as a weighted sum (*W_i_*) of its *m* nearest genes on both sides (*G_i_*). For genes at the ends of chromosomes, if there were less than *m* genes on one side, the number of remaining genes was used instead of *m* for that side.

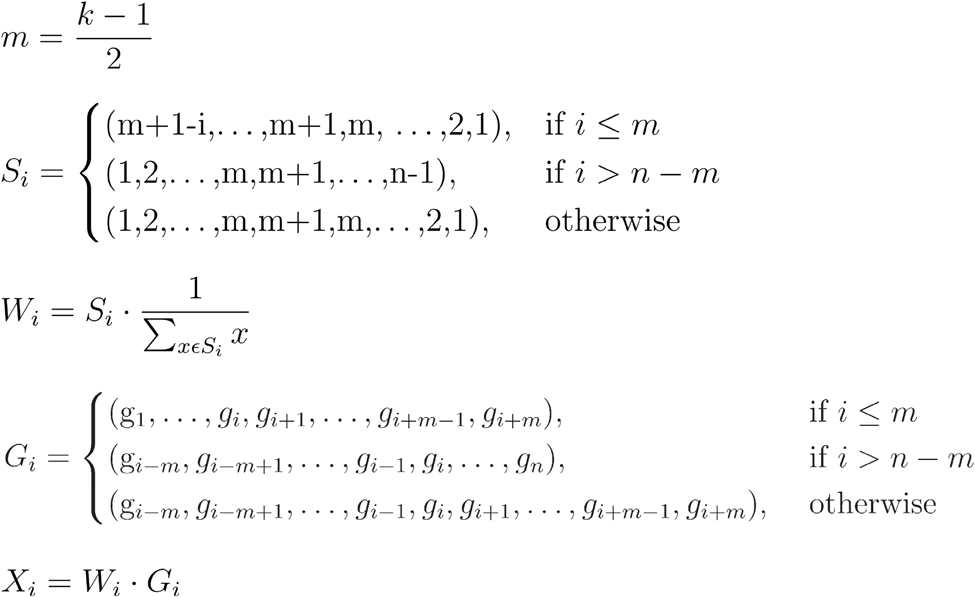

Each gene was re-centered by subtracting the mean-smoothed expression intensity of both malignant and non-malignant beads from each bead to create a baseline of zero expression intensity change. Then, the mean-smoothed expression intensity of reference beads was subtracted and the log2 transformation was reversed to rescale to expression intensities, resulting in a gene expression intensity matrix (beads x genes).

Following this expression smoothing, to address sparsity in Slide-Seq data, multiple adjacent beads were binned together, separately for non-malignant and malignant beads, under the assumption that spatially proximal and expression-similar beads are likely to have similar CNAs. To quantify bead spatial proximities, a bead spatial distance matrix **X**_*spatial*_, beads x beads) was calculated from the bead spatial matrix (beads x dimensions, where dimensions = 2 corresponding to the x and y axes) using Euclidean distance. To quantify how close beads are in expression space, the gene expression intensity distance matrix (**X**_*expression*_, beads x beads) was calculated from the gene expression intensity matrix (beads x genes) using Euclidean distance. Integrating these, beads were binned by pseudo-distance (**X**_*pseudo*_, beads x beads), defined as a linear combination of the bead spatial distance matrix **X**_*spatial*_, beads x beads) and the gene expression intensity distance matrix **X**_*expression*_, beads x beads). Both the distance matrix and relative gene expression intensity matrix were weighted by the adjustable parameters *k_spatial_* and *k_expression_*, respectively. *k_spatial_* = 55 and *k_expression_* = 1 were used here to prioritize binning beads that are spatially close together:

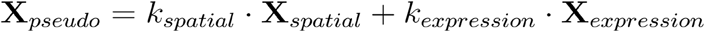

Bins were plotted across space for visual inspection. The resulting pseudo-distance matrix of all beads was hierarchically clustered using Ward’s hierarchical agglomerative clustering method (18) (ward.D2) with Euclidean distance, with the number of bins set to be the number of beads / 12 (**Supplementary Fig. 6d-f**). The expression score of each bin was calculated as the average of the relative gene expression intensities of beads assigned to that bin. Spatial coordinates of each bin were set to be the average x- and y-coordinates of its constituent beads. Clusters and type (malignant or non-malignant) of each bin were set to be the mode of those of their assigned beads. Lastly, the relative expression intensities in each bin were scaled by the number of UMIs per bin and expression values were capped at ≤ 1.4 and ≥ 0.6 to generate CNA scores.

### Clone determination

The malignant bins-only CNA scores (bins x genes) were hierarchically clustered by Ward’s hierarchical agglomerative clustering method(18) (ward.D2) with Euclidean distance with the cluster package v2.1.1 (19) in R and the number of clusters (putative clones) was selected by testing *k* = 2 to 10 clusters, and choosing the *k* that maximized the Silhouette score, *k*max. All non-malignant bins are assigned to one additional cluster.

### Differential gene expression analysis between CNA clusters

Differentially expressed genes were identified between a given CNA bead cluster and all other clusters (p_adj < 0.05) by a non-parametric Wilcoxon rank sum test (15) with the Bonferroni correction as implemented in Seurat v4, and ordered by decreasing average log-2 fold change. Genes were tested for enrichment in Gene Ontology 2018 Biological Processes using Enrichr (17,20).

### *In silico* Slide-Seq simulated data

To generate an *in silico* Slide-Seq dataset, nucleus profiles were annotated for cell types and assigned to pseudo-spatial coordinates, followed by Gaussian smoothing of expression values to obtain simulated Side-seq beads.

SnRNA-seq profiles for HTAPP-895-SMP-7359 and HTAPP-944-SMP-7479 were annotated as non-malignant or malignant based on SingleR v1.8.0 annotations (as described above) (16). A 3000×3000 square was created *in silico* to mimic the 3000 μm diameter of a Slide-seq puck. Two *in silico* spatial datasets were generated with either: (1) non-malignant nuclei spatially separate from malignant nuclei (“separated”) or (2) non-malignant nuclei mixed with malignant populations (“mixed”). In each scenario, 6,000 non-malignant nuclei profiles were sampled (with replacement) from the two snRNA-seq datasets combined (to provide distinct clones) and placed either at the top of the square at coordinate region [1:3000, 2701:3000] (“separated”) or across the square at coordinate region [1:3000, 1:3000] (“mixed”). Next, 7,000 malignant nuclei were randomly sampled (with replacement) from each sample. For the “separated” scenario, malignant nuclei profiles were placed at the bottom of the square at coordinate region [1:3000, 1:2700]; for the “mixed” scenario, malignant nuclei profiles were placed across the coordinate region [1:3000, 1:3000]. To test for recovery of clonal CNA profiles, malignant nuclei profiles were titrated from each sample to create a rightward gradient of malignant HTAPP-895-SMP-7359 profiles with decreasing frequency and a leftward gradient of malignant HTAPP-944-SMP-7479 profiles with decreasing frequency. These gradients were established by dividing the square into 100 zones of horizontal width = 30 so that zones had HTAPP-895-SMP-7359:HTAPP-944-SMP-7479 ratios in 1% decrements (99:0, 98:1, … 0:99), creating a square containing 20,000 nuclei (6,000 non-malignant, 7,000 malignant HTAPP-895-SMP-7359, and 7,000 malignant HTAPP-944-SMP-7479) across a 3000×3000 region with gradient mixture of the two malignant nuclei populations and either a separated or mixed non-malignant population.

Beads were simulated from the spatially positioned snRNA-seq profiles by spatial Gaussian smoothing across the 3000×3000 square, with a relative Gaussian kernel width = 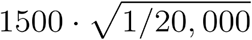 for 10,000 beads with randomly sampled x and y coordinates within the square and using actual bead size 10 μm (3) and cell size 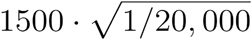 estimates. This created a set of 10,000 beads with two-dimensional spatial coordinates and combined expression values of constituent nuclei. The resulting *in silico* bead x gene expression matrix was used as input to SlideCNA and InferCNV for validation.

### Sensitivity analysis by down-sampling

For sensitivity analysis, reads from nucleus profiles used for simulation were downsampled to 25%, 10%, and 2% of their original counts across the entire dataset with the R package scuttle (21). Note that 10% downsampling led to a median of 867.5 and 647 reads per bead for the “separated” and “mixed” simulations, respectively, which is comparable to the median bead read count for real Slide-seq data (645 and 687 for HTAPP-895-SMP-7359 and HTAPP-944-SMP-7479 with reference transfer) after filtering to only keep beads with > 300 counts across all genes.

## Declarations

### Ethics approval and consent to participate

All samples used in this study underwent IRB review and approval at Dana Farber Cancer Institute (DFCI) (protocol 05-246) as well as the at the Broad Institute (protocol #15-370B).

### Consent for Participation

Not applicable

### Availability of data and materials

Code for SlideCNA (R package) is available through Github (https://github.com/dkzhang777/SlideCNA) and Slide-seq and snRNA-seq data are available as part of the HTAN-HTAPP data release on Synapse (syn20834712).

### Author contributions

J.K. designed the study with input from N.W. and A.R.. D.Z. developed SlideCNA and performed the validation analysis with guidance from J.K. and input from S.M.. A.S., and E.M. generated the Slide-seq data with input from F.C.. M.S. and J.W. generated snRNAseq data with input from O.R.R.. E.M., O.A., and J.K. processed snRNAseq and/or Slide-seq data. O.C. processed and analyzed exome sequencing data. D.A., J.J.-V., A.L., K.H., and S.R. assessed, archived, coordinated, and/or reviewed samples. D.Z., A.R., and J.K. wrote the manuscript with input from all co-authors.

## Acknowledgements

We thank all patients and their families. And we thank L. Gaffney for help with figure preparation.

## Author’s Information

Not applicable

---

**Fig. S1.**
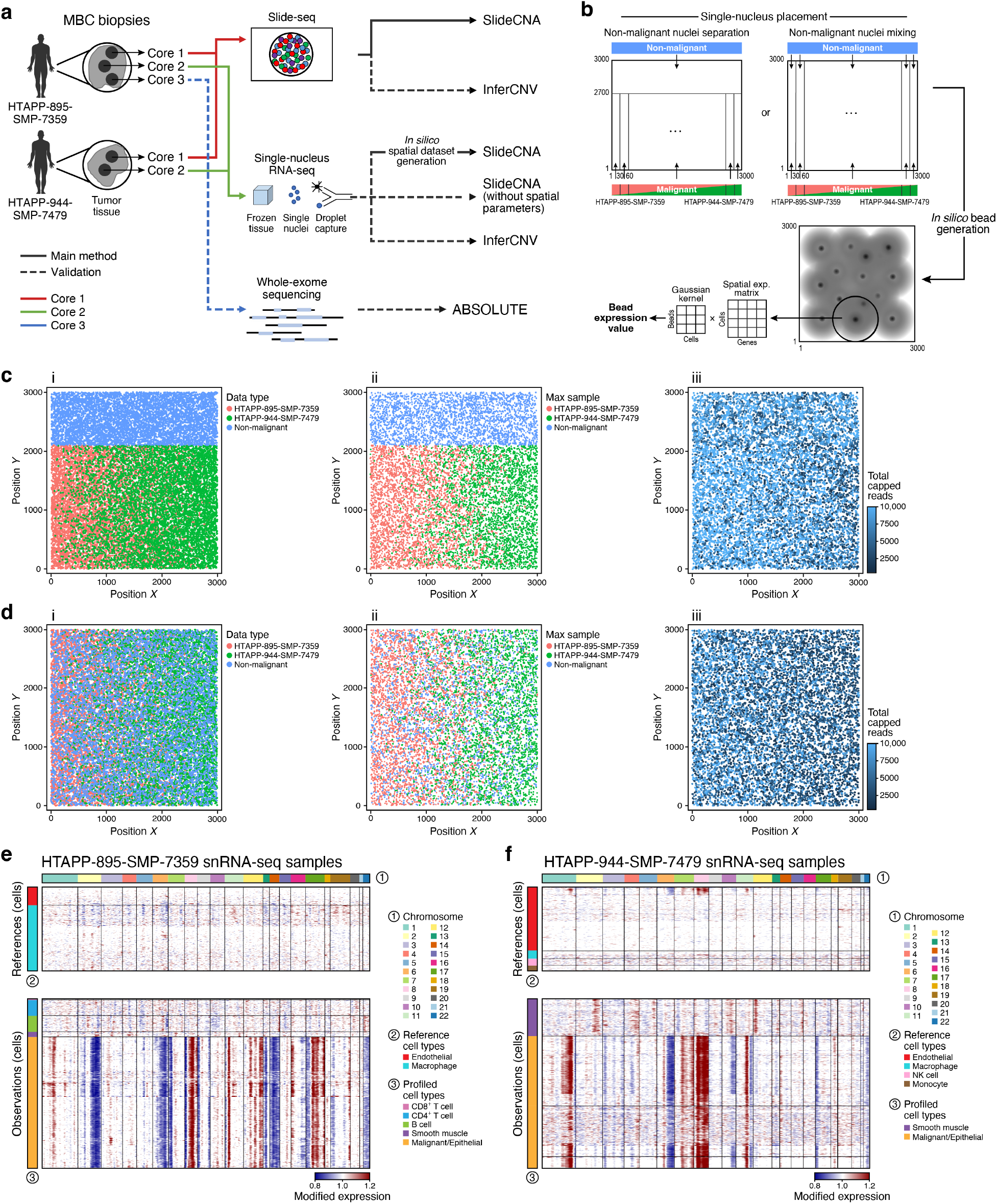
Workflow of SlideCNA implementation and validation, *in silico* schematic, and *in silico* dataset creation. **a,** High-level workflow of metastatic breast cancer data collection, Slide-seq and *in silico* spatial CNA analysis with SlideCNA, and validation with snRNA-seq and WES data. **b**, Schematic of generating the *in silico* datasets by placing snRNA-seq nuclei from each sample on a 3000×3000 square with either non-malignant nuclei separation or mixing and applying a Gaussian kernel to create *in silico* beads. **c,d,** i. Spatial plots of snRNA-seq nuclei; ii. the sample contributing the most reads to each *in silico* bead; and iii. the number of reads (capped at 10,000) per *in silico*-generated bead without downsampling for the *in silico* datasets with non-malignant separation (**c**) and non-malignant mixing (**d**). **e,f**, InferCNV heat maps of HTAPP-895-SMP-7359 and HTAPP-944-SMP-7479 snRNA-seq samples, respectively.

**Fig. S2.**
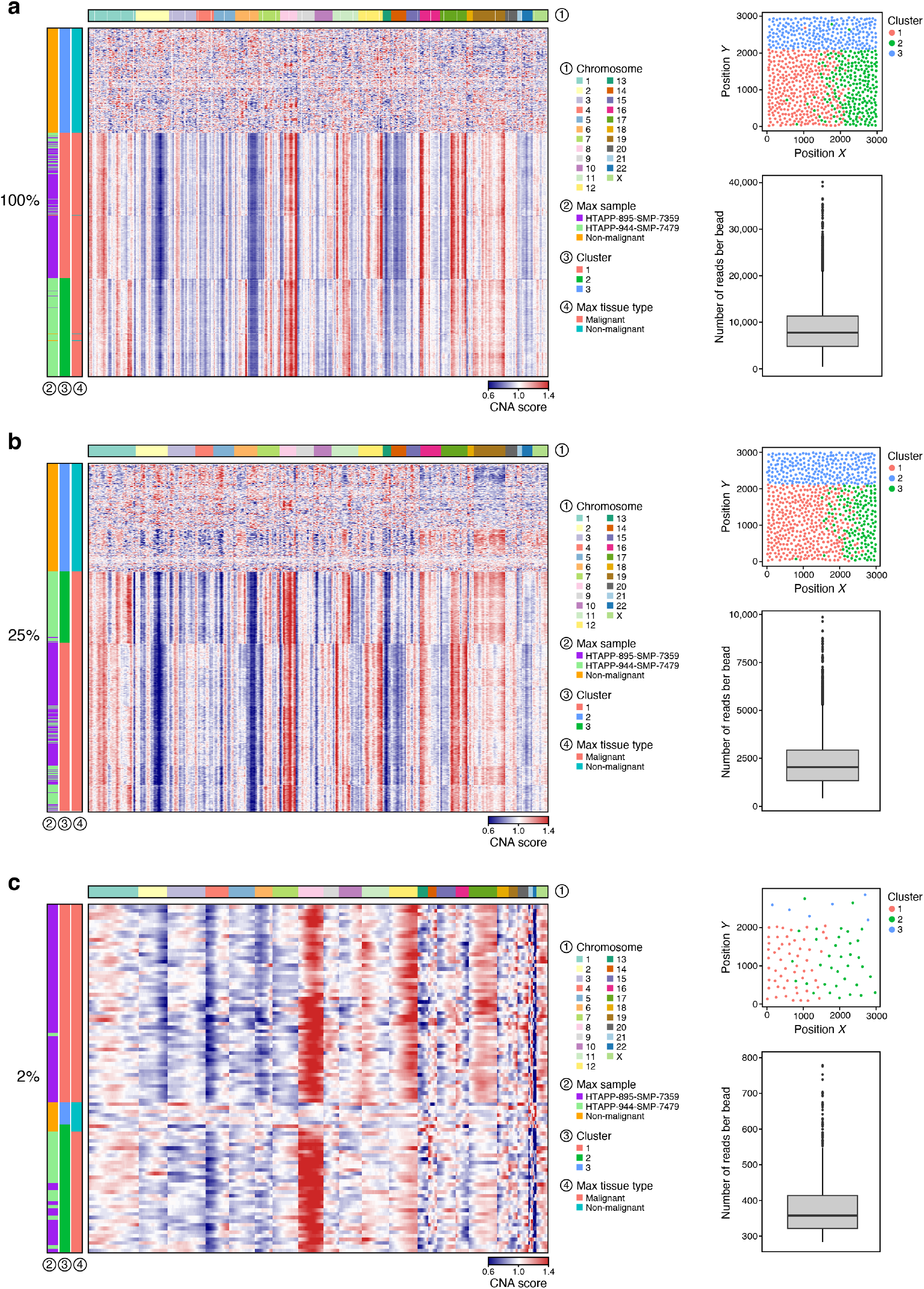
*In silico* spatial dataset with non-malignant separation results for downsampled counts. **a-c,** CNA heat map, spatial plot of bins colored by assigned cluster, and boxplot of number of reads per bin after filtering for beads with >300 counts across all genes for the *in silico* spatial dataset with non-malignant separation with varying degrees of downsampling: without downsampling (**a**), downsampled to 25% (**b**), and downsampled to 2% (**c**).

**Fig. S3.**
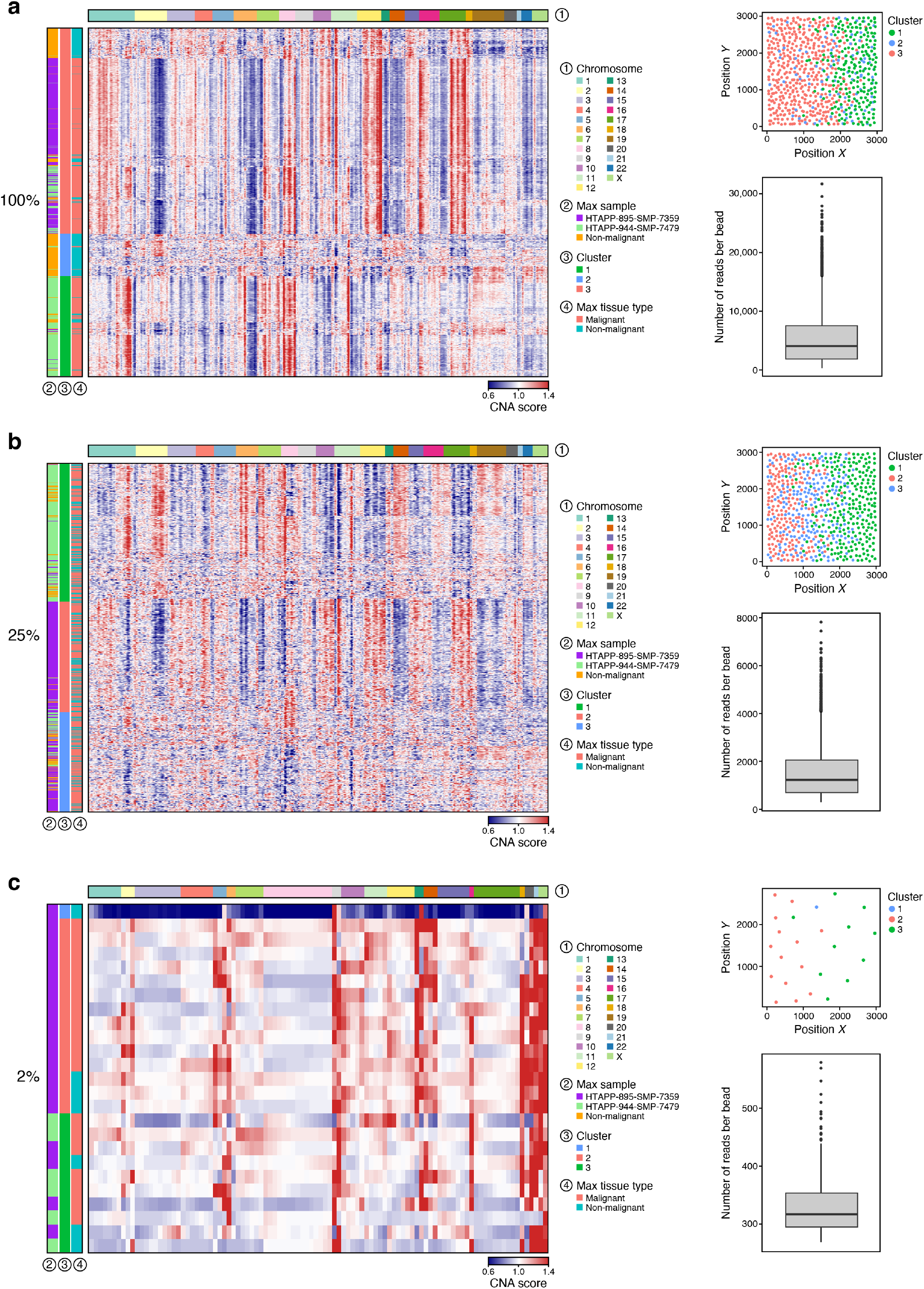
*In silico* spatial dataset with non-malignant mixing results for downsampled counts. **a-c,** CNA heat map, spatial plot of bins colored by assigned cluster, and boxplot of number of reads per bin after filtering for beads with >300 counts across all genes for the *in silico* spatial dataset with non-malignant mixing and counts with varying degrees of downsampling: without downsampling (**a**), downsampled to 25% (**b**), and downsampled to 2% (**c**).

**Fig. S4.**
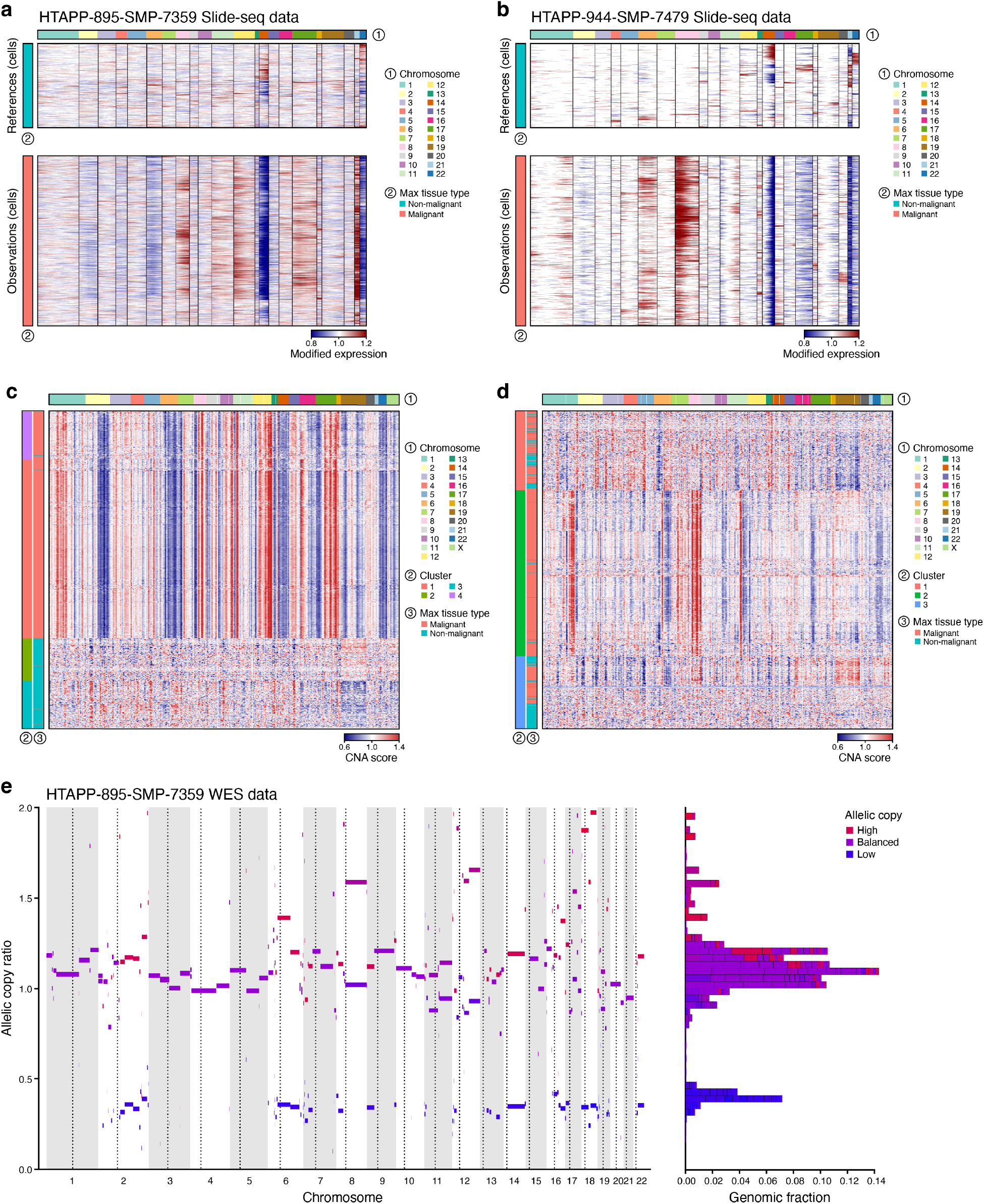
Slide-seq validation with InferCNV, snRNA-seq validation with SlideCNA, and WES CNA profile validation ABSOLUTE. **a,b,** InferCNV heat maps of HTAPP-895-SMP-7359 **(a)** and HTAPP-944-SMP-7479 **(b)** Slide-seq data, using the same reference beads as those determined for SlideCNA implementation. **c,d,** SlideCNA heat maps of HTAPP-895-SMP-7359 **(c)** and HTAPP-944-SMP-7479 **(d)** snRNA-seq data. **e,** ABSOLUTE CNA plot of the HTAPP-895-SMP-7359 WES data.

**Fig. S5.**
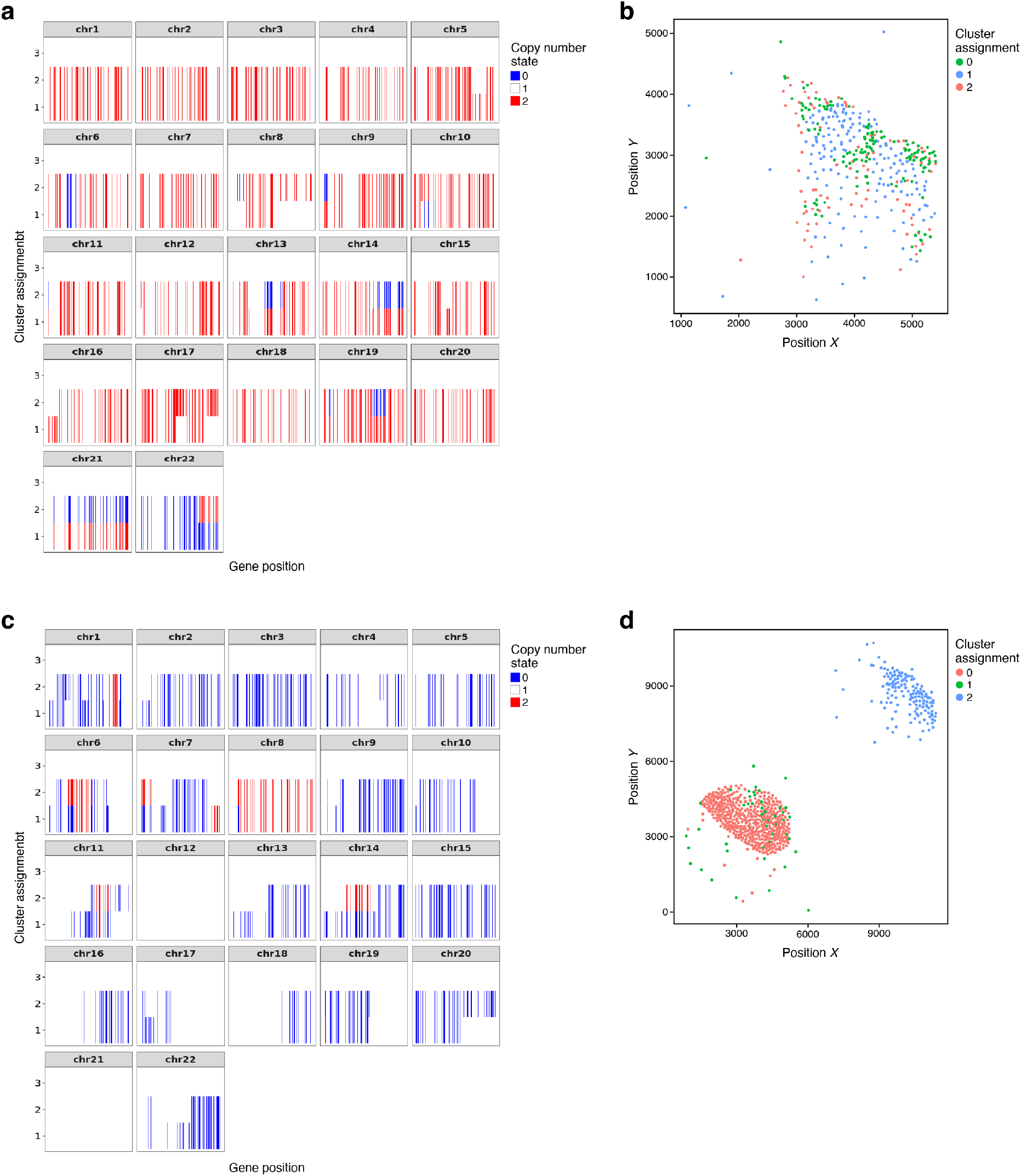
STARCH produces CNA profiles and cluster assignments. **a-d,** Copy number (CN) states assigned by STARCH per cluster **(a,c)** and spatial plots of STARCH-designated cluster assignments for binned beads **(b,d)** on Slide-seq data for HTAPP-895-SMP-7359 **(a,b)** and HTAPP-944-SMP-7479 **(c,d)**. Clusters 1 and 2 represent malignant binned beads and cluster 3 represents reference binned beads for both samples.

**Fig. S6.**
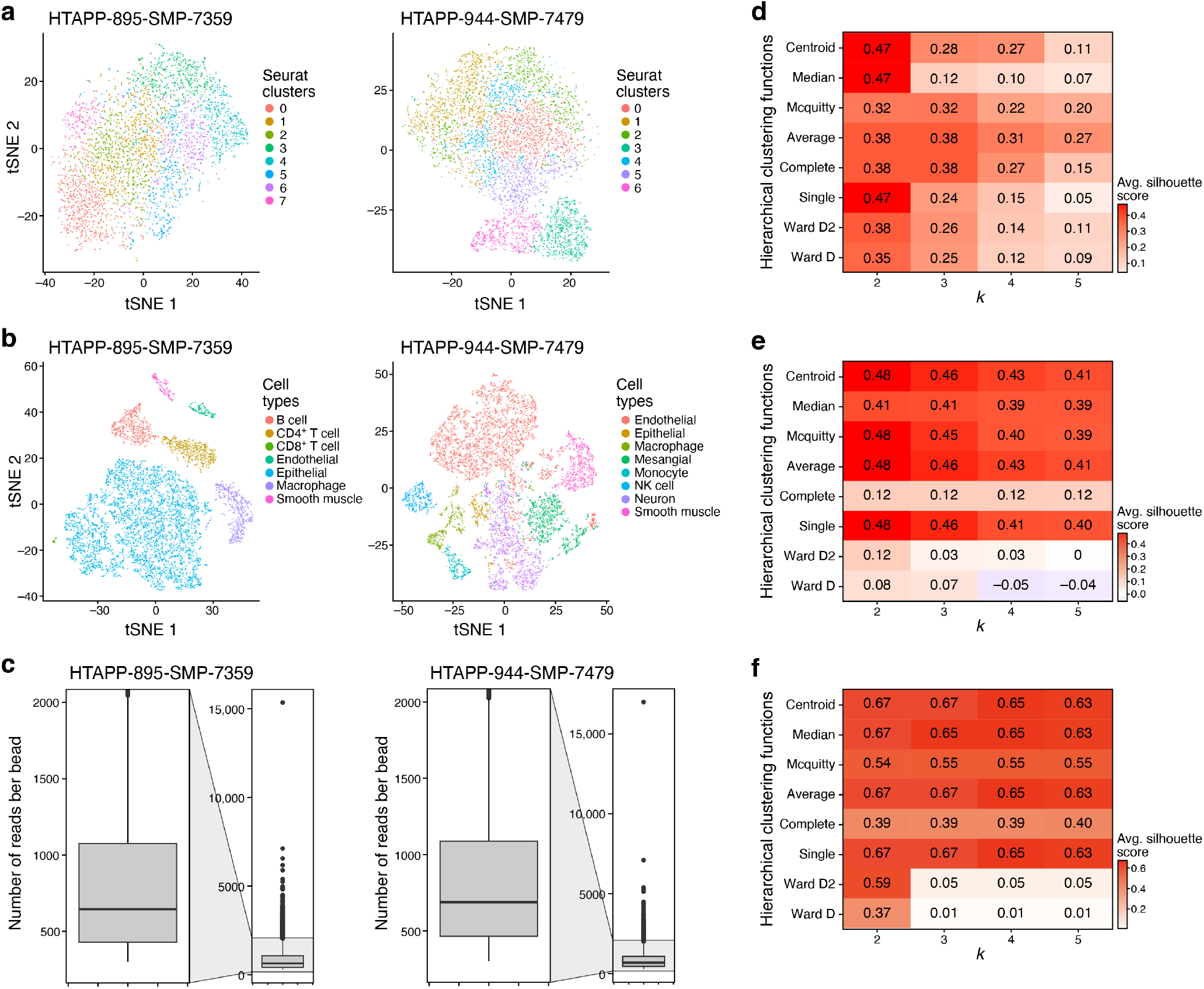
Sample embeddings, Slide-seq sample read distributions, and hierarchical clustering Silhouette scores. **a,b,** t-SNE plots of Slide-seq data Seurat clusters **(a)** and snRNA-seq data cell types **(b)** for the HTAPP-895-SMP-7359 and HTAPP-944-SMP-7479 samples. **c,** Boxplots of number of reads per bead after filtering for beads with > 300 counts across all genes for the two Slide-seq MBC samples. **d-f,** Silhouette scores for each hierarchical clustering method applied to determine the number of clusters, k, of malignant binned beads across k = 2 to 5 in the *in silico* dataset with non-malignant separation **(d)**, HTAPP-895-SMP-7359 Slide-seq sample **(e)**, and HTAPP-944-SMP-7479 Slide-seq sample **(f)**. A Silhouette score = 1 is the best, 0 is neutral, and −1 is poor.

